# A transcriptomic study of PF06830 root cap proteins

**DOI:** 10.1101/2024.01.29.577848

**Authors:** James A. Raymond

## Abstract

PF06830 is a family of about 2000 root cap proteins (RCPs) that are almost certainly involved in the major functions of the root cap, which include root growth and development, obtaining nutrients and sensing environmental variables. They appear to be expressed in the outer cell layers of the root tip where they are in intimate contact with the soil. Surprisingly, almost nothing is known about their individual functions, and they have received virtually no attention since their first description a quarter century ago. RCPs have easily identifiable characteristics and can be found in almost all plant species. The enormous expansion of plant transcriptomes in recent years provides an opportunity to better understand their functions, i.e., to see what biotic and abiotic variables affect their expressions. Here, the expressions of RCP genes in 49 root transcriptome studies (representing 14 species) obtained under different environmental conditions and at different development stages were investigated. (deleted sentence) In 19 of these studies, RCP expressions were found to be positively affected by environmental or developmental factors in specific cultivars of Arabidopsis, barley, rye, wheat, rice and cucumber. However, several negative effects were also found, often in different cultivars of the same species. These studies represent a first step in understanding the functions of RCPs that should help in the design of further studies. RCPs share some structural properties with, and may have overlapping functions with, other plant protein families, including small heat shock proteins, late embryogenesis abundant proteins and lectins. Their origins appear to predate the development of roots.

## Introduction

The roots of almost all land plants have a layer of cells called the root cap or calyptra that protect the root tip and allow it to communicate with the surrounding soil (Kumpf & Nowack, 2015). Its functions were a source of wonder to Charles Darwin a century and a half ago (Darwin & Darwin, 1880). The structure and function of the root cap cell layer have been the subject of many studies (Arnaud *et al*., 2010, Barlow, 2002, Berhin *et al*., 2019, Iijima *et al*., 2008, Rüger *et al*., 2023, Sievers *et al*., 2002). Surprisingly, however, none of these studies examined a large family of proteins (PF06830) that are specifically expressed in the root cap. The first mention of proteins in this family was in 1999, in which two ∼320-a.a. proteins were reported to be specifically expressed in the root caps of maize (Matsuyama *et al*., 1999). Searching the relatively meager resources in GenBank at that time, they found two genes (one in white spruce and one in Arabidopsis) that matched the C-terminal ∼250-a.a. portions of the maize proteins. Since then, it appears that only one other study has mentioned this domain: it was found in a freshwater alga and used to identify it as a streptophyte (Nedelcu *et al*., 2006). For reasons that are not clear, NCBI’s conserved domain database recognizes only a ∼60-a.a. region at the C-terminus of these proteins, calling it the root cap domain and assigning it to protein family PF06830. Interestingly, the PFAM database (now incorporated into InterPro) has 2000 proteins listed in this family, all of which have sizes similar to those reported by Matsuyama et al. as well as the C-terminal ∼60-a.a. root cap domain. Interpro also pipelined these proteins into AlphaFold to predict their structures. Surprisingly, no analyses of their structures or functions could be found. In the following, PF06830 proteins are referred to as root cap proteins (RCPs).

Recently, my colleagues and I described an ice-binding protein from a streptophyte glacier alga (*Ancylonema nordenskioeldii*; *An*IBP) that consists of two domains, a DUF3494 domain that is well known to bind to ice, and a 250-a.a. C-terminal domain that was clearly a member of the PF06830 (RCP) family (Procházková *et al*., 2024). AlphaFold showed that the C-terminal domain was largely composed of a β-sandwich, in which two parallel β sheets form a sandwich-like structure, followed by a ∼60-a.a. root cap domain, whose structure is not known. A blast search of GenBank using the 250-a.a. C-terminal domain of *An*IBP as a query yielded about 2000 proteins, in agreement with the number of proteins listed in PF06830. All of the hits were streptophyte proteins. An analysis of randomly selected proteins from this group using RNA-seq data from NCBI showed that they were almost entirely expressed in root tissue (Procházková et al., 2024). RCPs consist of three domains: a variable N-terminal domain that starts with a secretion signal followed by a conserved domain encoding a β-sandwich structure and the 60-aa. root cap domain (Fig. 1). The N-terminal domains are typically short (∼100 a.a.) regions with unknown functions. AnIBP is the only known exception, in which this region serves as the ice-binding domain. In unicellular streptophytes, the start of the conserved domain (i.e., the start of the β-sandwich domain) begins with DPH (Asp-Phe-Hist), and in land plants it begins with DPR (Asp-Phe-Arg) with few exceptions.

**Fig. 1.**
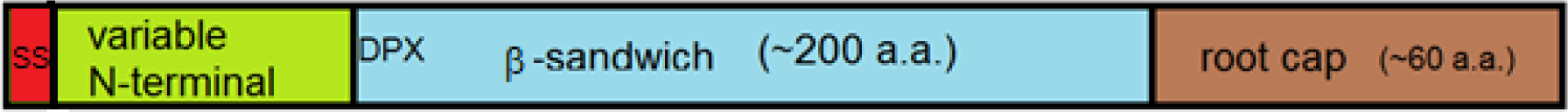
Structure of root cap proteins. The N-terminal domain is usually around 100 a.a and usually begins with a secretory signal (SS). The β-sandwich domain begins with DPH (unicellular streptophytes) or DPR (land plants). The root cap domain is a conserved domain; an attempt to predict its structure with AlphaFold was unsuccessful (Procházková et al., 2024).

Some grasses have dozens of RCPs and a streptophyte alga (*Closterium*) has over 50 of them (Procházková et al., 2024). Interestingly, RCPs share several properties with small heat shock proteins (sHSPs), including high abundance and diversity, small size, typically <25 kDa, a central region encoding a β-sandwich structure and a variable N-terminal domain (Waters and Vierling 2020). sHSPs in plants as well as other organisms are upregulated in response to abiotic stresses and have been proposed to act as chaperones that prevent damage to proteins by such stresses (Haslbeck & Vierling, 2015, Waters & Vierling, 2020). This raised the possibility that RCPs also serve to mitigate abiotic stresses. Such a hypothesis is supported by the finding that the root cap is intimately involved in sensing and responding to changes in the soil environment (Kumpf & Nowack, 2015). To test this idea, we previously looked at RNA-seq data in NCBI that examined the effect of cold stress on the *Triticum aestivum* (wheat) root transcriptome (Procházková et al., 2024). We found that the expressions of three of its RCP genes went through some cyclic changes as the temperature was lowered from 18 °C to 2 °C, confirming they were responsive to cold stress.

As a first step in understanding the functions of RCPs, the goal of the present study was to see what factors influence their expressions. For this, I used RNA-seq data deposited in NCBI, which has more than 8000 studies (BioProjects) on the effects of various biotic and abiotic factors on streptophyte transcriptomes. As might be expected, most of these studies focused on economically important crops. The present study focused on root transcriptomes because previous studies (Matsuyama et al., 1999, Procházková et al., 2024) showed that RCPs were almost exclusively expressed in root tissue. Forty-nine of these studies that focused on root transcriptomes were selected for the present study. The results show that many RCPs respond to stress factors and changes in development.

## Methods

Previous work (Procházková et al., 2024) showed that RCPs were almost entirely expressed in root tissue. To obtain root transcriptomes, Google Scholar and the NCBI SRA database were searched for key words (root, transcriptome, RNA-seq, stress) and various stress terms such as cold, heat, drought, etc. The search yielded a total of 49 NCBI BioProjects. For each species, typically multiple homologous RCPs were found with NCBI protein blast searches using the conserved domains of existing RCPs as queries. For rye and sugarcane, for which no RCP homologs were found in GenBank, hundreds of RCP transcripts were obtained by blastn searches and assembled into 200-nt queries by Geneious. When the number of RCPs for a given species was large, sequences that appeared to be representative of the group were selected. For each RCP, a 200-nt-long sequence starting with the DPR motif was selected as a query. The queries were then used in online blastn searches of the BioProject’s transcriptomes using the parameters max. no. hits =5000 and e-value = <1e-10. When two RCP queries got the same hit, the hit was assigned to the RCP that had the lowest e-value. The number of hits in each transcriptome was expressed as number per million reads in the transcriptome. Since the query lengths were 200 nt (0.2 kb), the FPKM (fragments per kilobase of transcript per million mapped reads) value can be obtained by multiplying this number by 5 (1/0.2).

All data were processed and plotted with Excel (Microsoft). Signal peptides were identified with SignalP 6.0 (https://services.healthtech.dtu.dk/services/SignalP-6.0/). Conserved domains were identified with NCBI’s conserved domains tool (https://www.ncbi.nlm.nih.gov/Structure/cdd/wrpsb.cgi). A maximum likelihood phylogenetic tree was made with Mega X (Tamura *et al*., 2021).

## Results

A total of 67 RCPs were used as transcriptome queries in this study (Table S1). Full-length sequences are available for 61 of them in GenBank. All but two of these were confirmed to have two characteristics identified in the N-terminal region of maize RCPs by Matsuyama et al., a secretory signal and multiple pairs of cysteine residues. This appears to be the first confirmation of these properties in other members of the PF06830 family. In addition, all of the full-length RCPs were confirmed to have a ∼60-a.a. root cap domain near their C-termini as shown by NCBI’s conserved domain database.

Forty-nine BioProjects that examined root transcriptomes were screened for RCP expression (Table S2). Expressions of the RCPs were highly variable, often varying among themselves in a given species, among cultivars of the same species and among different species, or not showing any effect (Table S2). Nineteen of the studies in which either positive or negative changes in RCP expression were observed are described here.

### Arabidopsis thaliana

BioProject PRJNA793162 (Pacheco *et al*., 2022) obtained root transcriptomes for six Arabidopsis ecotypes (representing different root hair phenotypes) that had been subjected to cold treatment (3 days at 10 °C vs. 22 °C control). *A. thaliana* has at least six RCPs. Five of them were selected as representative (Table S2). Four did not show clear changes but one of them (AT1) appeared to be up-regulated by cold treatment in all six ecotypes (Fig. 2A).

**Fig. 2.**
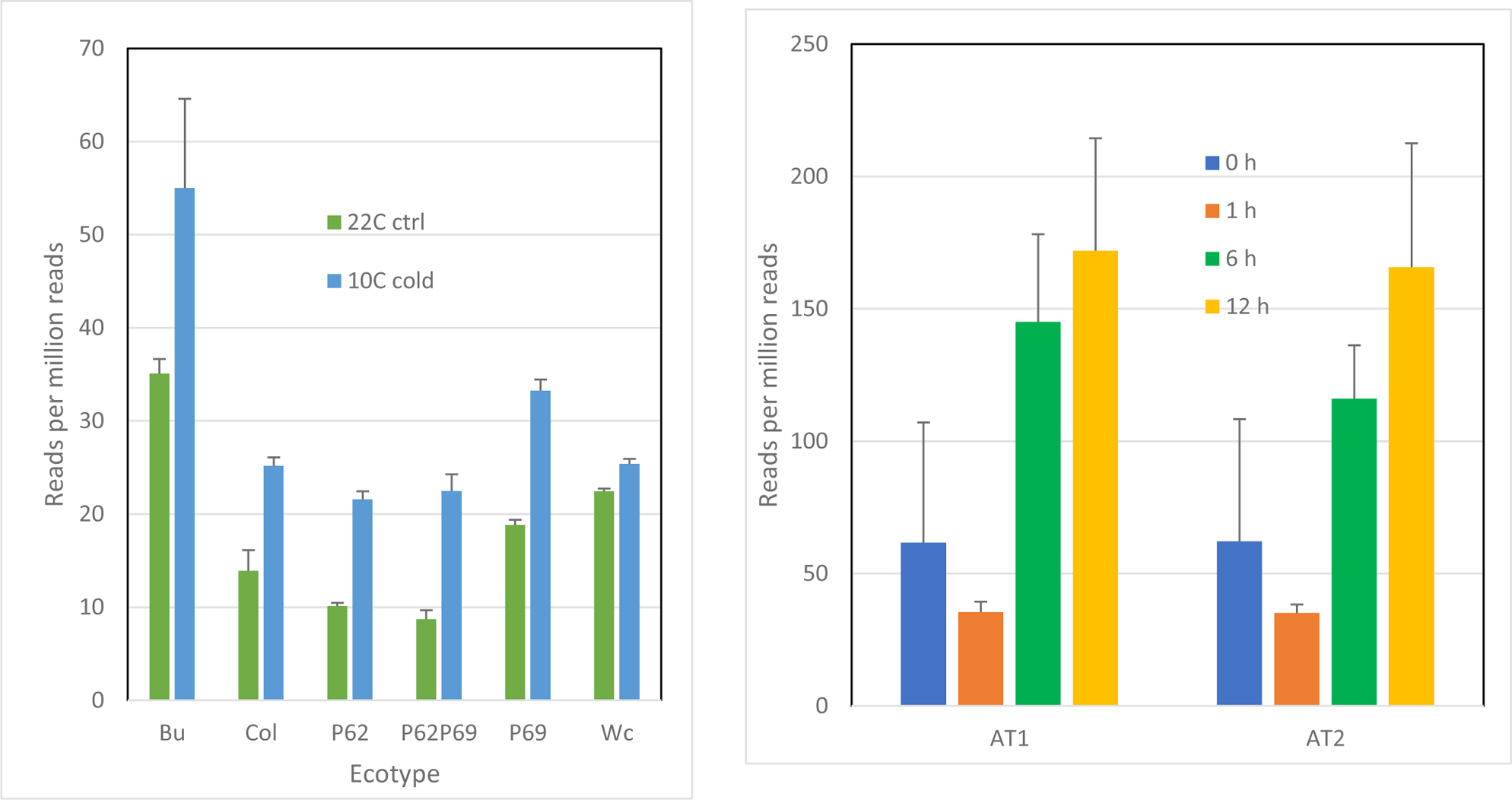
Effects of stresses on the expression of a RCP genes in *Arabidopsis thaliana*. A. Effect of cold stress on AT1 expression in six *Arabidopsis* ecotypes. Error bars show range of two transcriptome replicates. B, Effect of lead exposure time on expression of two RCP genes in *Arabidopsis* wild type (Col-0). Error bars show standard deviations of three transcriptome replicates.

Bioproject PRJNA593333 (Zheng *et al*., 2021) examined the effect of lead (1 mM) for different exposure times on gene expression in Arabidopsis (Col-0). Three of the five RCP genes were only modestly affected by lead exposure, while two of them (AT1 and AT2) appeared to be upregulated as the time of exposure increased from 0 to 12 h (Fig. 2B).

### Hordeum vulgare (barley)

BioProject PRJNA896249 (Wang *et al*., 2023) examined the effects of low N treatment (0.1 mM nitrate for 3 and 18 days) on the root transcriptomes of two barley cultivars. Barley has about 14 RCPs. Three of them (HV1-HV3, Table 1) were selected as representative. Treating W20, a low N-sensitive cultivar, to low N conditions increased the expressions of all three RCP genes for both treatment periods (Fig. 3A), while treating W26, a low N-tolerant cultivar to the same conditions had the opposite effect, significantly decreasing their expressions (Fig. 3B).

**Fig. 3.**
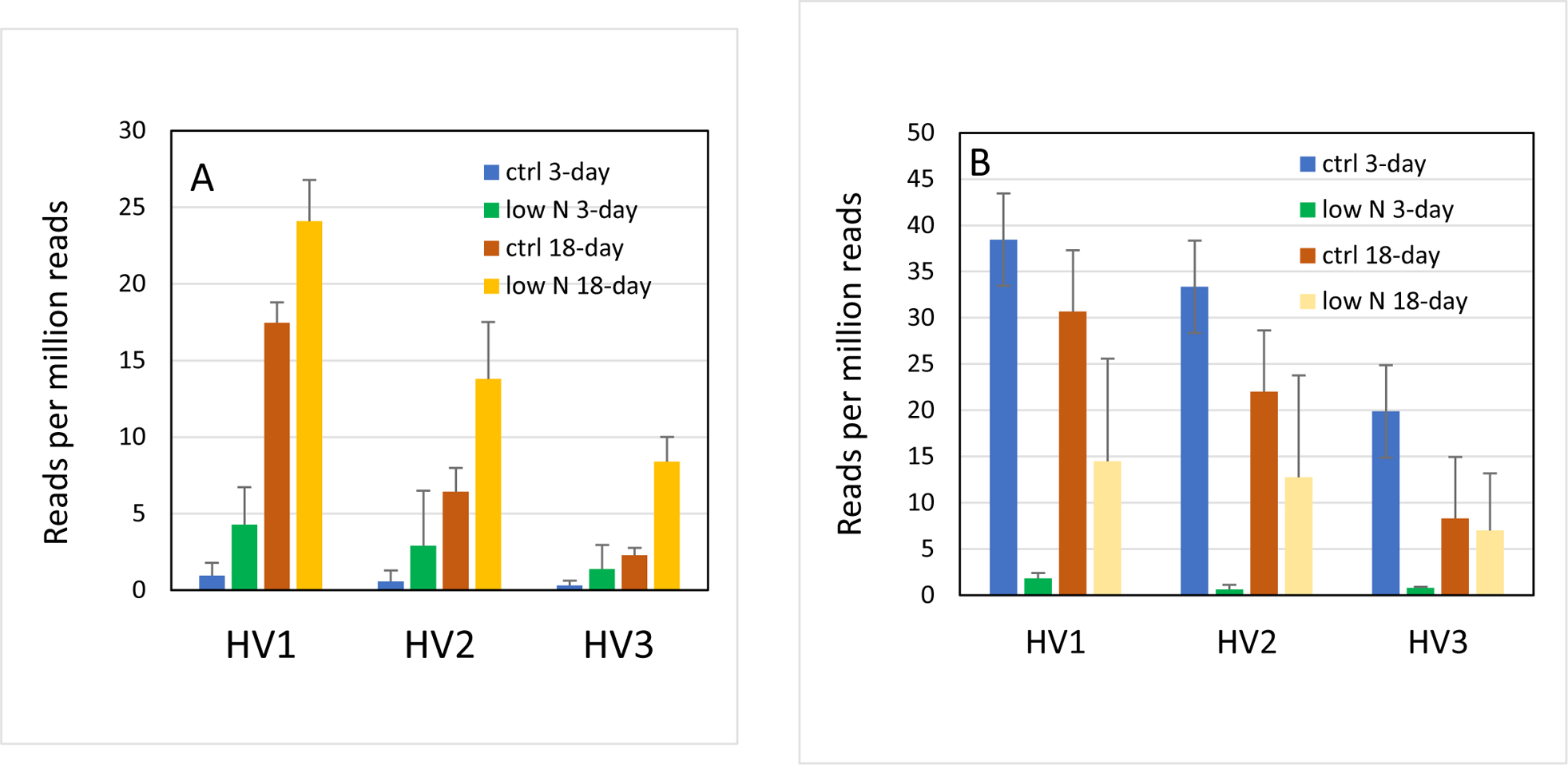
Effect of low nitrogen for 3 and 18 days on the expression of three RCP genes in two barley cultivars. A, cv, W20 (nitrogen sensitive). B, cv. W26 (nitrogen efficient). Error bars show the standard deviation of three transcriptome replicates.

### Triticum aestivum (wheat)

A blast search shows that the *T. aestivum* genome (BioProject PRJNA764258) has about 50 RCPs. For experiments with this species, nine of them were selected as representative.

Several studies have examined the effect of salt on salt-tolerant cultivars of *T. aestivum*, yielding conflicting results. In one experiment (PRJNA970414) (Yang *et al*., 2008), a salt-tolerant wheat cultivar (xiaobingmai 33) was subjected to salt treatment (150 mM Na+) for 15 days. This treatment down-regulated all nine RCPs in the root transcriptome (Fig. 4A). On the other hand, in two other salt-tolerant cultivars, RCP expressions were increased by salt treatment. One of them (PRJNA340343/−344) (Karam *et al*., 2022), examined the effects of long-term salt treatment (250 mM NaCl) on cv. Kharchia local plants. All nine wheat RCPs were upregulated in this experiment (Fig. 4B).

**Fig. 4.**
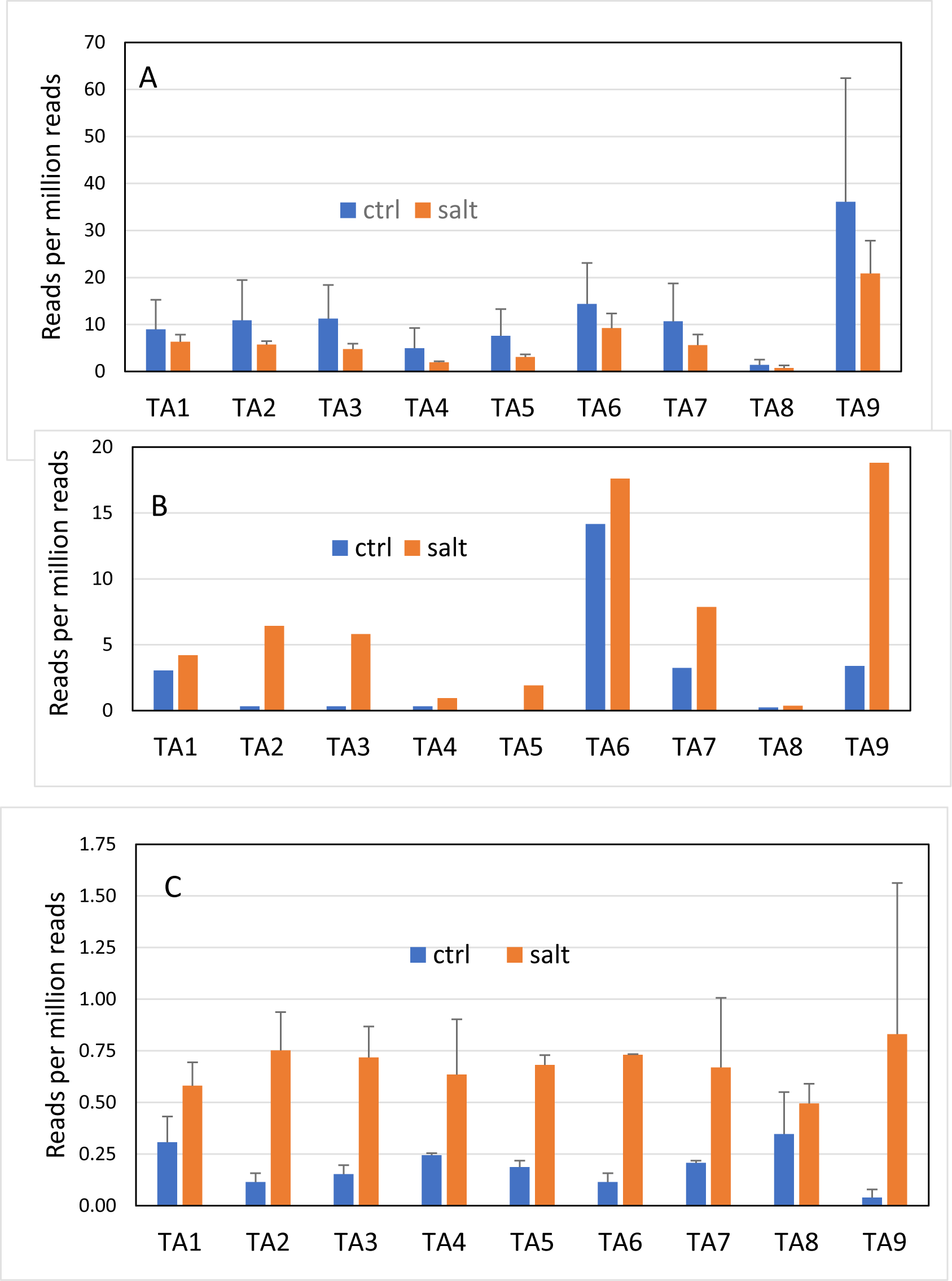
Effects of salt stress on the expression of nine RCP genes in the root transcriptomes of three salt-tolerant cultivars of *T. aestivum*. A, xiaobingmai 33. Error bars show the standard deviation of three transcriptome replicates. B, Kharchia local. C, arg. Error bars show the range of two transcriptome replicates.

In the other study (PRJNA487923) (Amirbakhtiar *et al*., 2019), 3-week old seedlings of the salt tolerant cultivar arg were treated to 150 mM NaCl for 12 h. RCP expressions were an order of magnitude less than they were with the xiaobingmai 33 and Kharchia local cultivars but they still were strongly upregulated by salt treatment (Fig. 4C). Another salt-tolerant wheat cultivar (Najran) (PRJNA936261) (Alyahya & Taybi, 2023) showed that salt treatment (200 mM NaCl for 4 weeks) had little effect on the expressions of any of the wheat RCPs (data not shown).

Soil microorganisms have important roles in shaping the diversity and productivity of land plants as well as providing protection against a diversity of stresses (Begum *et al*., 2019). BioProject PRJNA551129 (Campos *et al*., 2019) used a mycotrophic grass (*Lolium perenne*) to establish a healthy mycorrhizae population in the soil and then removed the grass. They showed that breaking up the hyphae (by sieving the soil) before planting wheat (cv. Ardila) had negative effects on growth and tolerance of Mn toxicity. In the case of the RCPs, breaking up the hyphae modestly upregulated their expressions in 1-week-old roots (Fig. 5), as if they were compensating for the loss of mycorrhizae. However, after five weeks of growth, the RCPs were all slightly down-regulated (data not shown), perhaps as a result of the roots adapting to the soil. Similar RCP expressions were obtained from the transcriptomes when another mycotrophic plant (*Ornithopus compressus*) was used to establish soil mycorrhizae (data not shown).

**Fig. 5.**
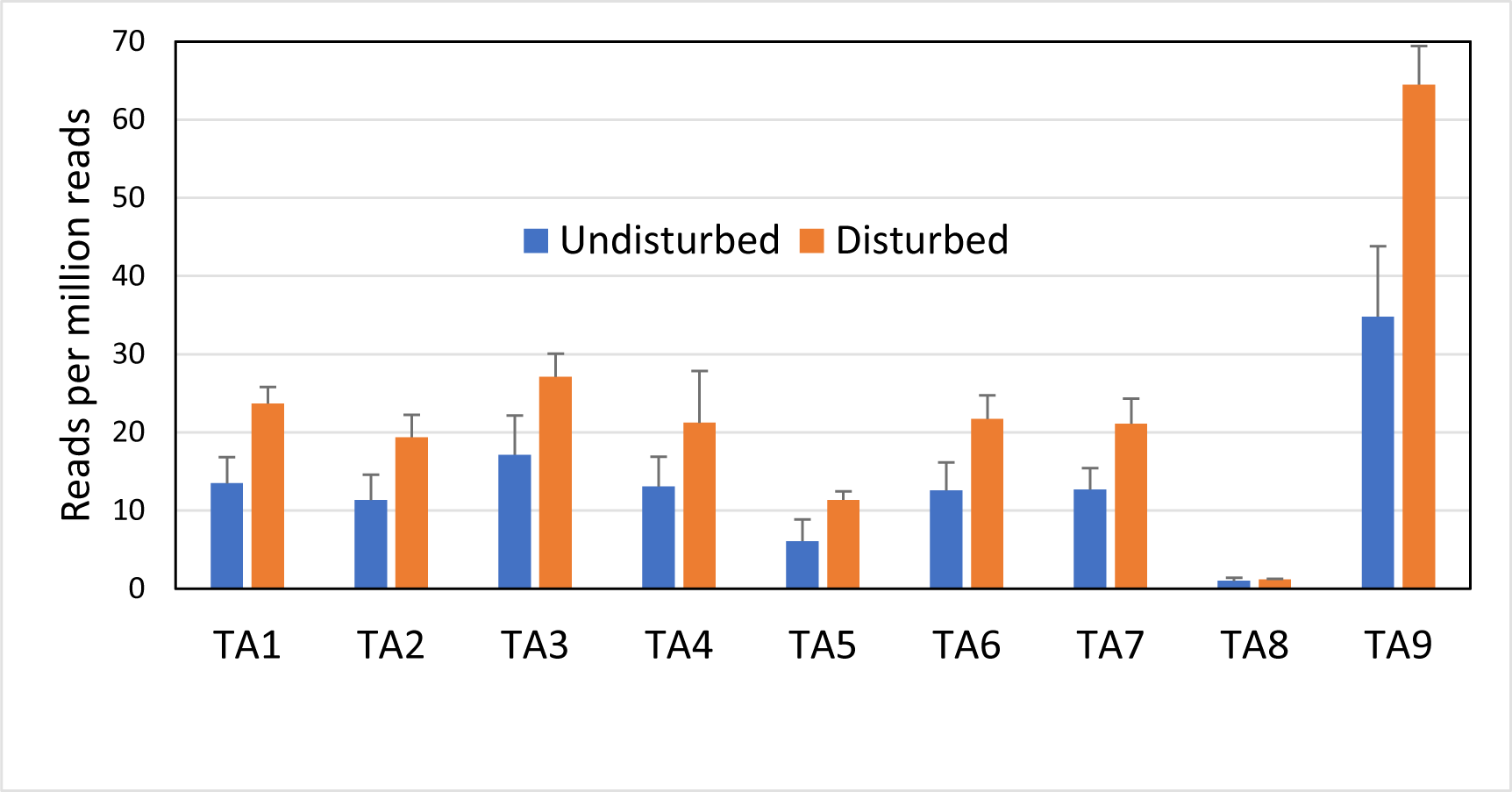
Effect of breaking up (disturbing) soil hyphae on the expression of nine *Triticum* RCP genes in 1-week-old wheat (cv. Ardila) roots. The mycorrhizae were established by first growing a mycotrophic grass (*Lolium rigidum*) in the soil and then removing the grass. Error bars show the range of two transcriptome replicates.

### Secale cereale (rye)

PRJNA564622 (Rabanus-Wallace *et al*., 2021) obtained root transcriptomes of *T. aestivum* and rye as the plants were cooled from 18 °C to 2 °C over a 70-day period. We previously reported that *T. aestivum* RCP genes peaked at least twice as the temperature was cooled (Procházková et al., 2024). Here, we examined their data for rye. RCP homologs could not be found in NCBI so three of them were assembled from the transcriptomes (Table 1). They were nearly identical to RCPs in *T. aestivum* over the 200-nt query range. As was observed with the *T. aestivum* RCPs, expression of the rye RCP genes was low but cyclic, with peaks at 3, 14, 42 and 63 days during the cooling period (Fig. 6).

**Fig. 6.**
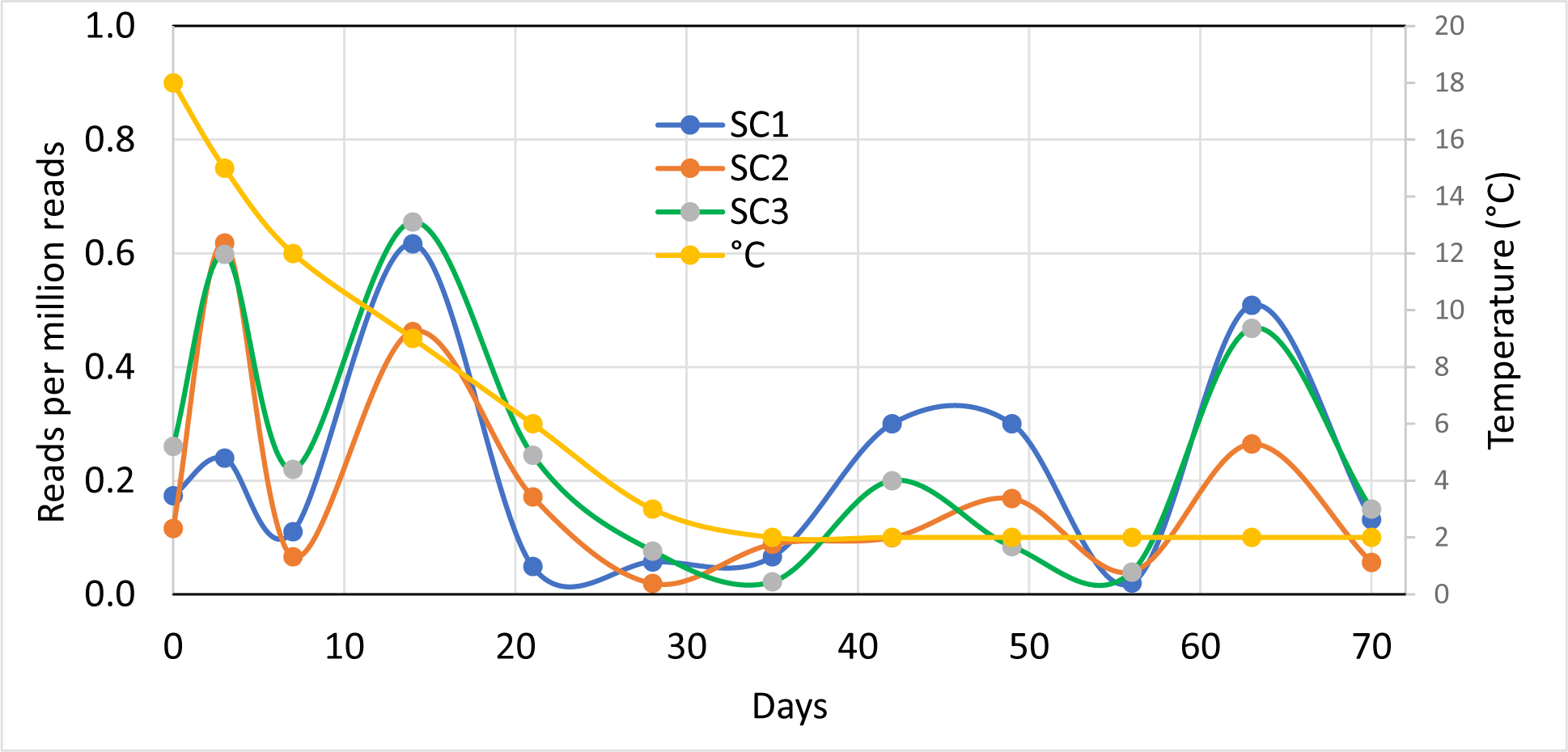
Expressions of three RCPs of *Secale cereale* during cold adaptation over a 70-day period.

### Zea mays (maize)

Maize has about six RCPs, four of which were selected as representative (Table 1). One of them (ZM1) is the same as ZMRCP1 described by Matsuyama et al. (1999).

To understand the mechanisms of salt tolerance in maize, BioProject PRJNA647980 (Luo *et al*., 2021) compared the transcriptomes of salt-sensitive and salt-tolerant cultivars after 0, 3, 6 and 18 h of salt stress (150 mM Na+). RCP expressions did not significantly change with increasing salt exposure, but they were generally downregulated in the salt-tolerant cultivar. The zero-hour (control) data are typical and are shown in Fig. 7A.

**Fig. 7.**
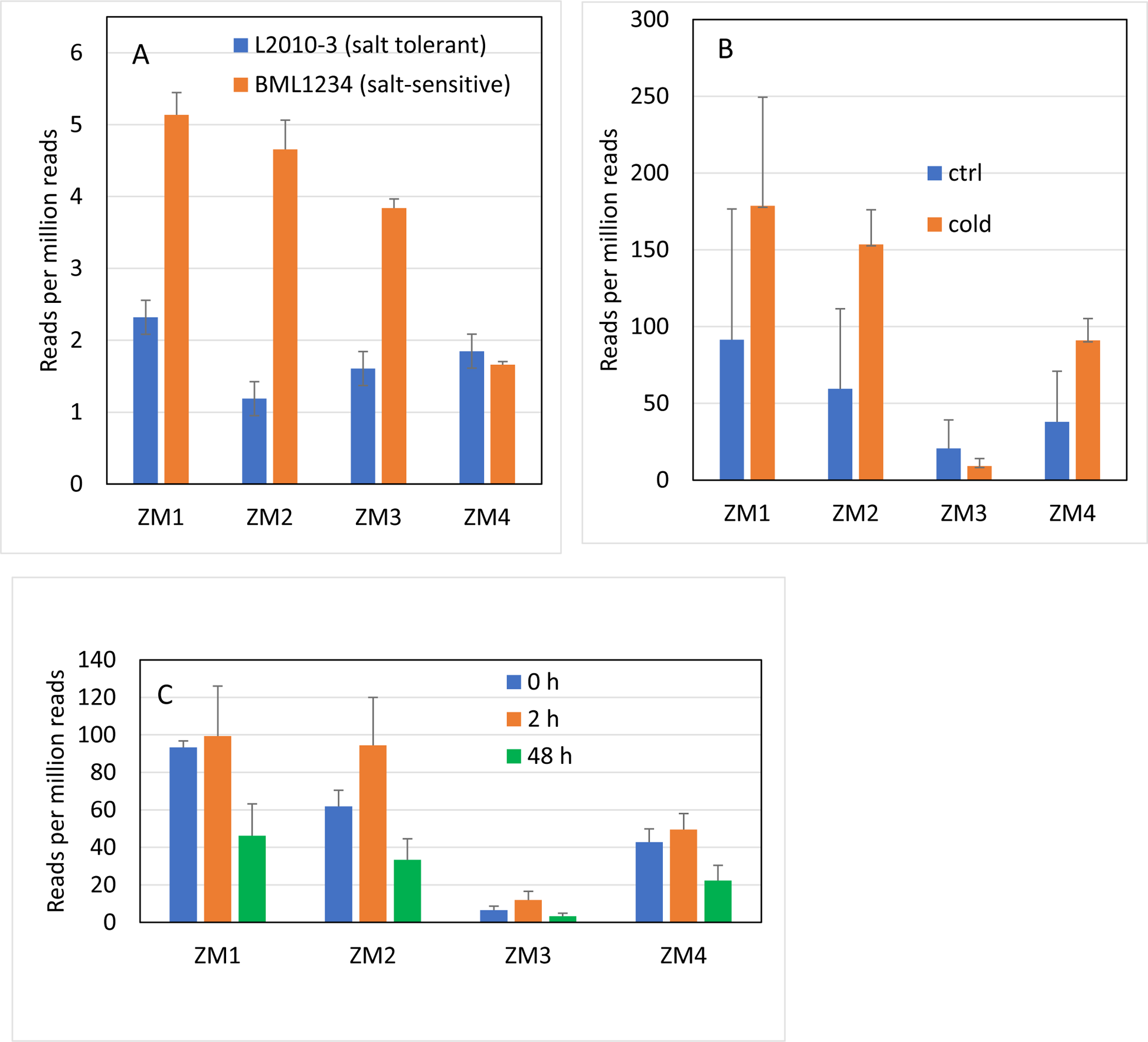
Expression of four RCPs in different maize cultivars. A, Comparison of salt-tolerant and salt-sensitive cultivars in the absence of salt stress. The patterns are roughly similar to those at 3, 6 and 18 h of salt exposure. B. Effect of cold exposure in maize cv. He344. C, Effect of 2 and 48 h heat stress on cv. B73. Error bars in each panel show the standard deviation of three transcriptome replicates.

BioProject PRJNA686250 (Zhao et al. 2021) looked for clues to the cold response mechanisms in the roots of cv. He344, a major maize variety grown in Northeast China. Two-week-old seedlings were exposed to 5 °C or 22 °C (control) for 3 days. ZM1, 2 and 4 of He344 were unusual in that their expressions were among the highest of the RCP expressions observed in this study (Fig. 7B). These three RCPs also appeared to be upregulated by cold stress.

BioProject PRJNA520822 (He *et al*., 2019) examined heat stress responses in maize by exposing cv. B73 to 38 °C for up to 48 h. As in cv. He344, RCP gene expressions were generally high. Two hours of heat stress had no clear effect on their expressions, while 48 hours of heat stress appeared to decrease their expressions by half (Fig. 7C).

### Saccharum spontaneum (sugarcane)

To better understand the mechanisms of cold resistance in sugarcane, BioProject PRJNA319158 (Dharshini *et al*., 2020) obtained root transcriptomes of a high altitude cultivar (IND 00–1037) after plants initially grown at 28 °C were exposed to 10 °C for up to 48 h. Three RCPs were assembled from the transcriptomes using wheat RCPs as queries. Two of them (SS1 and SS2) were strongly expressed before cold treatment and then their expressions gradually decreased with increasing cold exposure, while the relatively weakly expressed SS3 was not greatly affected by cold (Fig. 8).

**Fig. 8.**
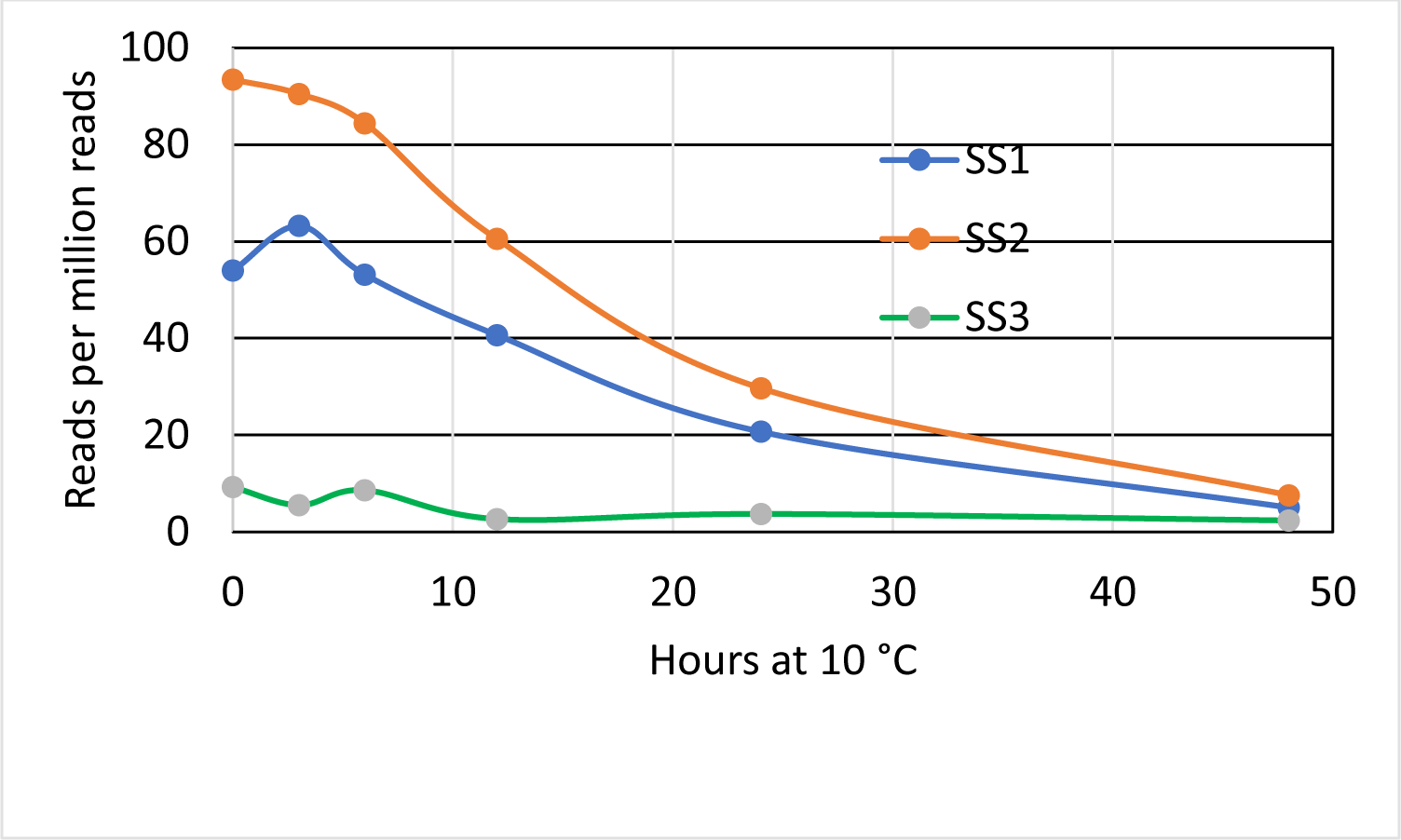
Effect of increasing exposure to cold stress on the expressions of three RCPs in sugarcane.

### Oryza sativa (rice)

Rice of the Japonica and Indica groups each have at least seven RCPs. For each group, five were selected as representative: OSj1-5 for the Japonica group and OSi1-5 for the Indica group (Table 1). Each of the five OSj genes was found to be homologous to one of the five OSi genes, so they were given the same numbers: OSj1 was homologous to OSi1 and so on.

BioProject PRJNA359218 (Yoo *et al*., 2017) examined the effect of two and three days of drought on gene expression in a 4-week-old japonica rice cultivar (Chilbo). All five RCP genes were strongly down-regulated (some were undetectable) by 2 and 3 days of drought (Fig. 9).

**Fig. 9.**
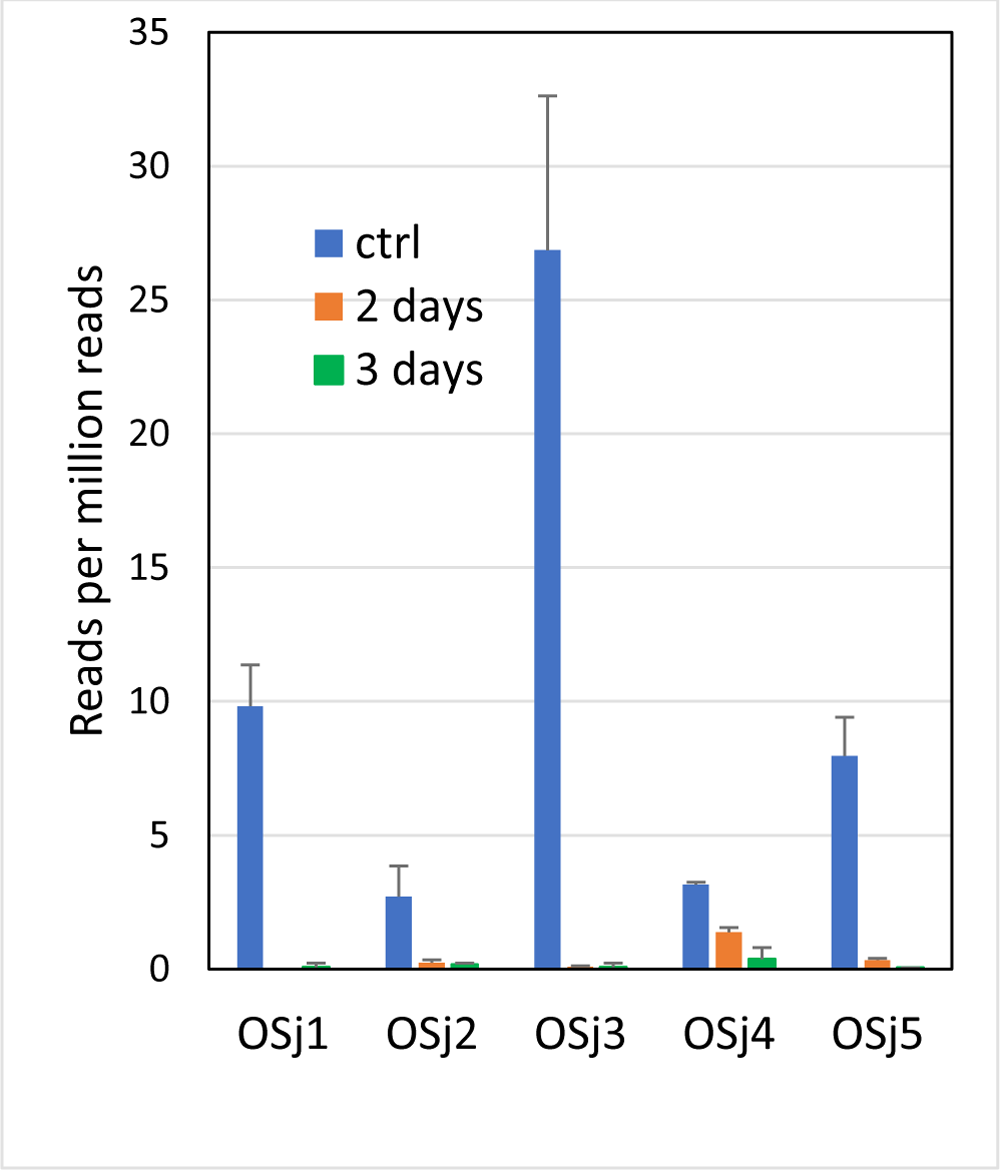
Effects of two and three days of drought on the expression of five RCP genes in the root transcriptomes of rice (cv. Chilbo). Few or no reads were found in most of the 2- and 3-day transcriptomes. Error bars show the range of two transcriptome replicates.

Bioproject PRJNA812782 (Deng *et al*., 2022) examined the effect of salt stress (140 mM NaCl for 3 h) on salt-tolerant and wild type cultivars of japonica rice to find genes involved in salt tolerance. In the wild-type (Fig. 10A), salt stress upregulated four of the RCP genes, one of them significantly, and downregulated one. Similar changes were observed in the salt-tolerant mutant (Fig. 10B).

**Fig. 10.**
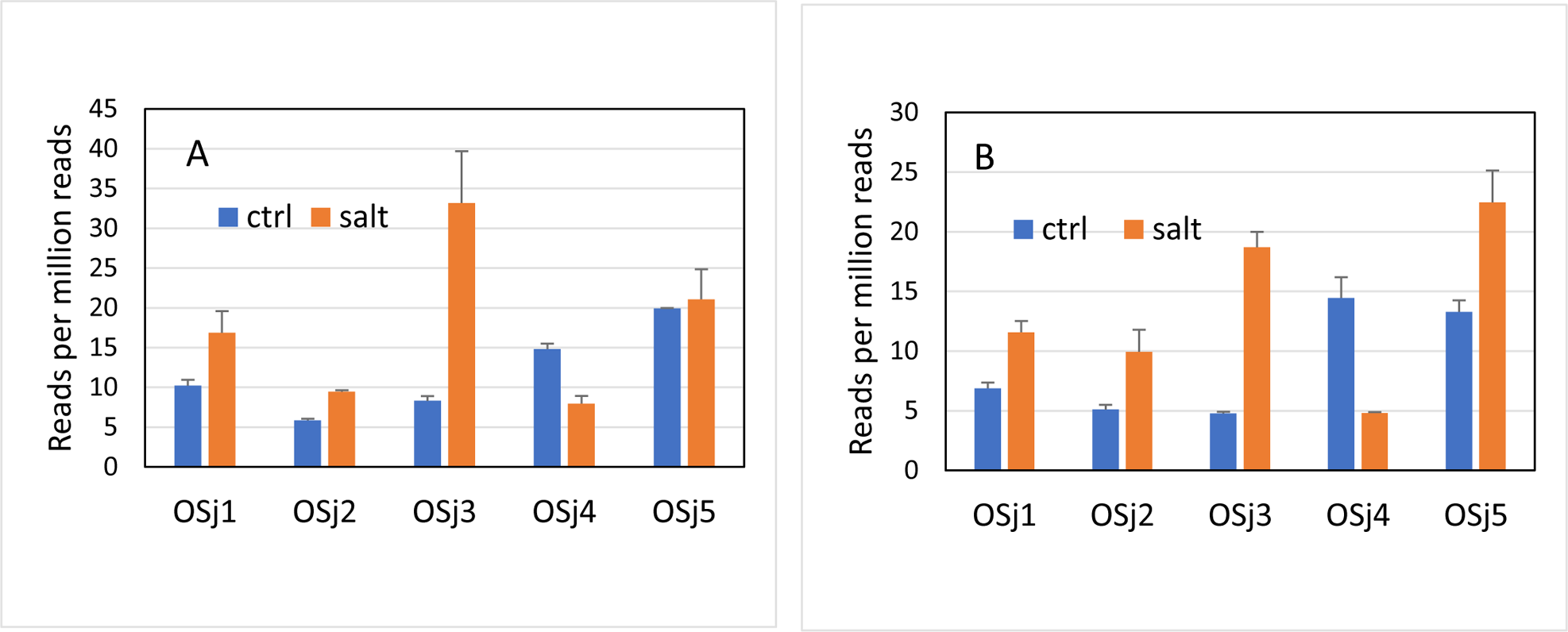
Effects of salt stress on the expressions of five RCP genes in the root transcriptomes of wild-type (A) and salt-tolerant (B) varieties of japonica rice. Error bars show the range of two transcriptome replicates (control) or the standard deviation of three transcriptome replicates (salt).

Rice has three types of root, lateral, nodal and S-lateral, with different sensitivities to salt stress. To understand the reasons for the different sensitivities, BioProjects PRJNA319757 and PRJNA342794 (Cartagena *et al*., 2021) examined the effect of salt stress (50 mM NaCl for 7 days) on the transcriptomes of the three root types in wild-type (IR29) and salt-tolerant (Mulai) japonica rice cultivars. Salt stress appeared to upregulate most of the RCP genes in both the lateral and nodal roots of both cultivars (Fig. 11), the exceptions being OSi3-5 in the wild-type nodal roots, which were down-regulated or unchanged by salt stress. Salt stress had little on RCP expressions in S-lateral roots (data not shown).

**Fig. 11.**
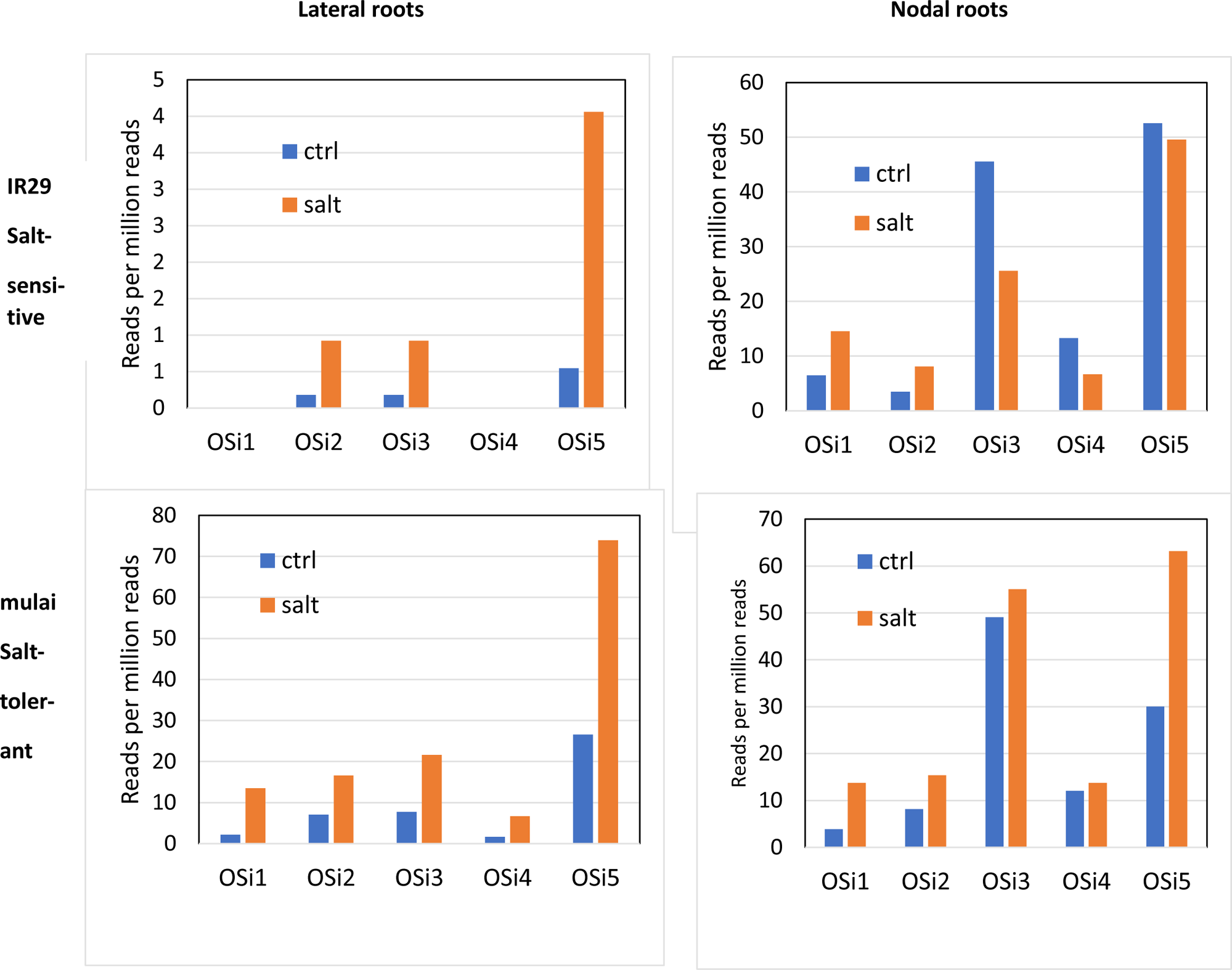
Effects of salt stress on the expressions of five RCP genes in the lateral root and nodal root transcriptomes of wild-type (IR29) and salt-tolerant (mulai) varieties of indica group rice.

Certain species of nematodes severely impact the growth of rice by feeding on their roots. Bioproject PRJNA589993 (Zhou *et al*., 2020) obtained root transcriptomes of a japonica group rice cultivar (Nipponbare) 6 and 18 days after the soil was inoculated with nematodes (*Meloidogyne incognita*) to see how the plants were responding to their attacks. Nematode infestation downregulated all five of the RCP genes after both 6 and 18 days of infestation (Fig. 12).

**Fig. 12.**
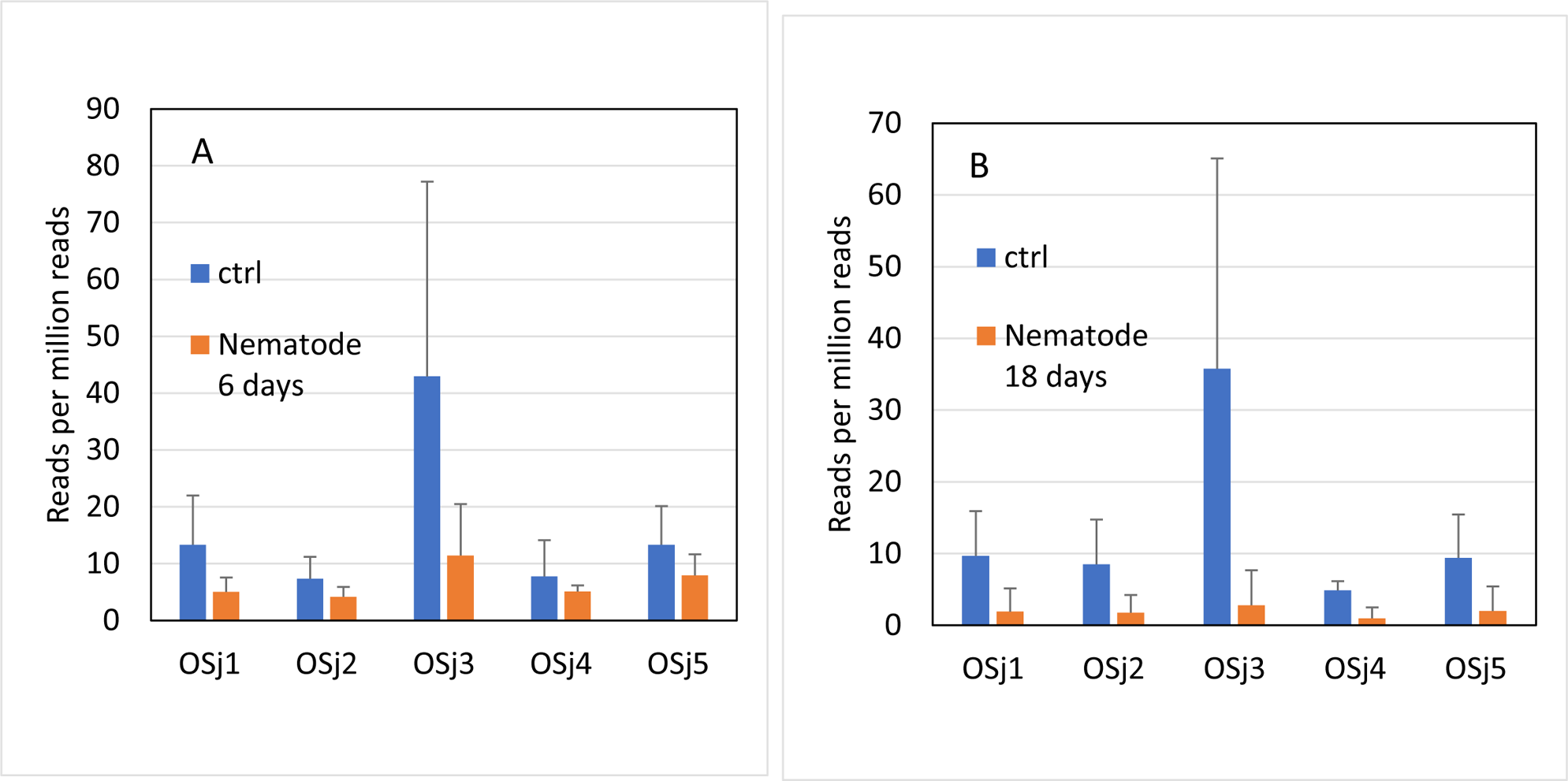
Effects of 6 days (A) and 18 days (B) of nematode infestation on the expressions of five RCP genes in rice (cv. Nipponbare). Error bars indicate standard deviation of three transcriptome replicates.

Bioproject PRJNA613213 (Sevanthi *et al*., 2021) attempted to identify quantitative trait loci (QTLs) associated with drought tolerance and low N tolerance in rice. To this end, they compared the root transcriptomes of two indica group cultivars from different habitats: IR64, which is adapted to lowland (irrigated) habitats, and Nagina 22, which is adapted to drier, upland habitats. IR64 is sensitive to both drought and low N, while Nagina22 is more tolerant of drought and low N. For drought stress, the plants were given only 20% of the normal watering for 7 days, while for low N stress, the plants were grown in soil containing only 1% of the normal N for 21 days. In both cultivars, OSi5 was the most strongly expressed RCP under all conditions (Fig. 13). In the lowland IR64, OSi5 was strongly upregulated by both low N and drought, while in Nagina s22, it was downregulated by low N and not strongly affected by drought. The other RCPs were either moderately upregulated or unaffected by the two conditions in both cultivars.

**Fig. 13.**
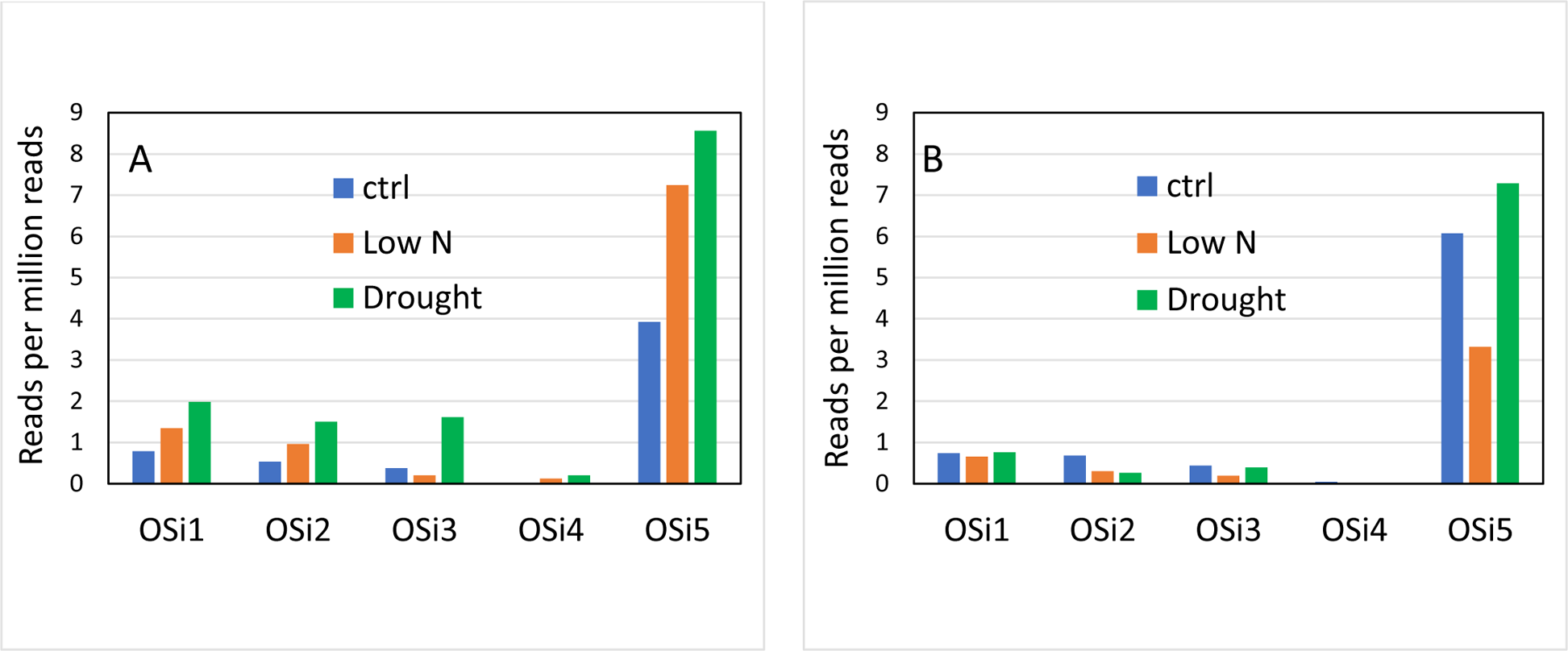
Effects of low N and drought on the expressions of five RCP genes in rice. A, IR64, a low N- and drought-sensitive cultivar. B, Nagina 22, a low N- and drought-tolerant cultivar. One transcriptome was obtained for each condition.

Arbuscular mycorrhizal fungi (AMFs) can have positive effects on plant growth, including the mitigation of salt stress in rice (e.g., (Tisarum *et al*., 2020)). To understand their effect on salt tolerance, Bioproject PRJNA827100 (Hsieh et al. 2022) examined the effects of *Rhizophagus irregularis* and salt stress (3 weeks exposure to 150 mM NaCl) on the root transcriptome of the japonica rice Nipponbare. In the present analysis, both AMF and NaCl tended to downregulate the five RCP genes, and their negative effects appeared to be additive (Fig. 14).

**Fig. 14.**
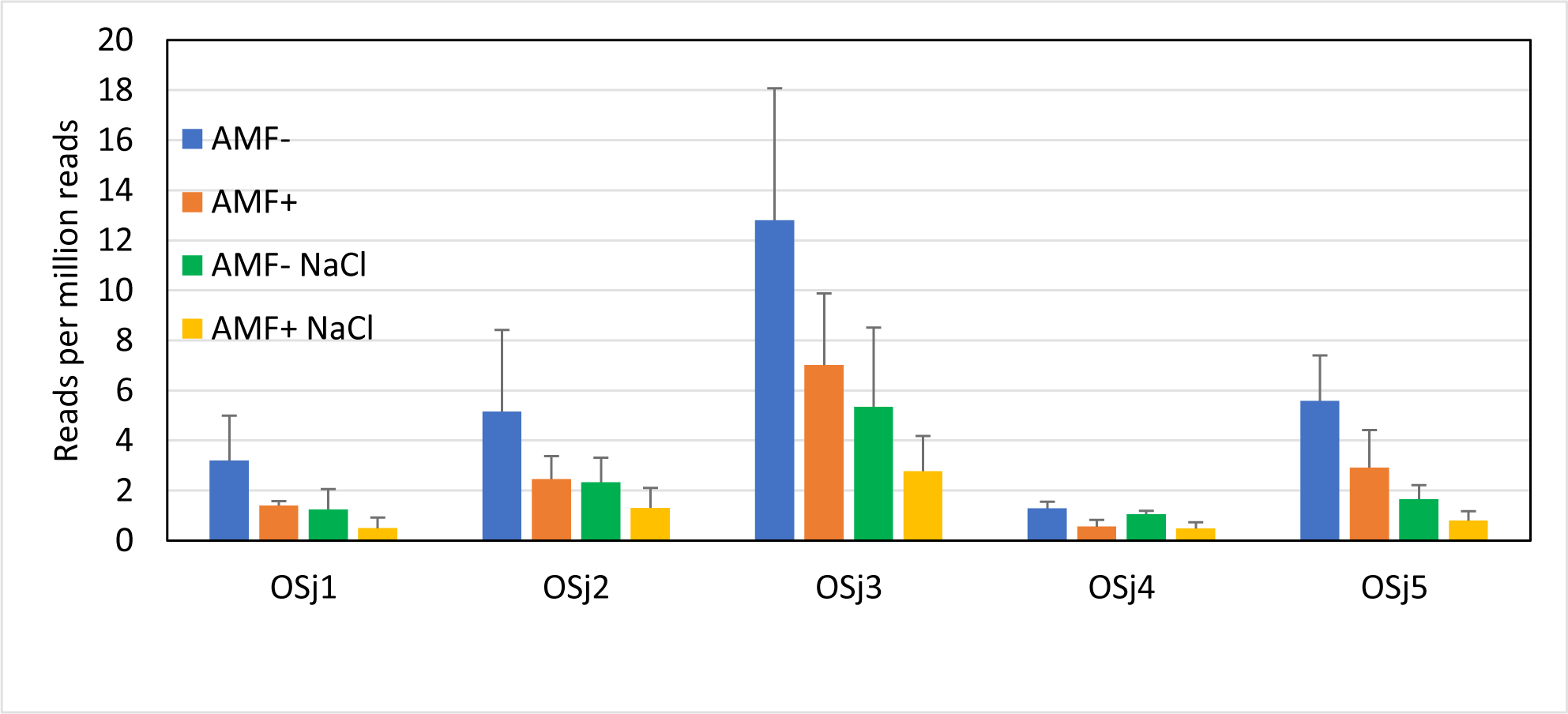
Effects of an arbuscular mycorrhizal fungus (*R. irregularis*) and NaCl (150 mM) on the expressions of five RCP genes in rice cultivar Nipponbare. Error bars show the standard deviation of three transcriptome replicates, except for the AMF+ bars (orange), in which they show the range of two transcriptome replicates.

### Cajanus cajan (chickpea) and Phaseolus vulgaris (common bean)

BioProjects PRJNA354681(Pazhamala *et al*., 2017) and PRJNA221782 (Chakraborty *et al*., 2023) obtained the root transcriptomes of chickpea (cv. ICPL 87119) and common bean (cv. Bat93), respectively, each at four stages of development. Chickpea has about 10 RCPs, five of which were selected as representative. Four of the five RCPs were expressed almost exclusively at the early (seedling and vegetative) stages and were almost undetectable at the later (reproductive and senescence) stages (Fig. 15A). *P. vulgaris* has 4 RCPs, only 3 of which were expressed in the transcriptomes. Each of them was also expressed more strongly in the early stages (6 and 12 days of growth) than in the later stages (29 and 43 days) (Fig. 15B).

**Fig. 15.**
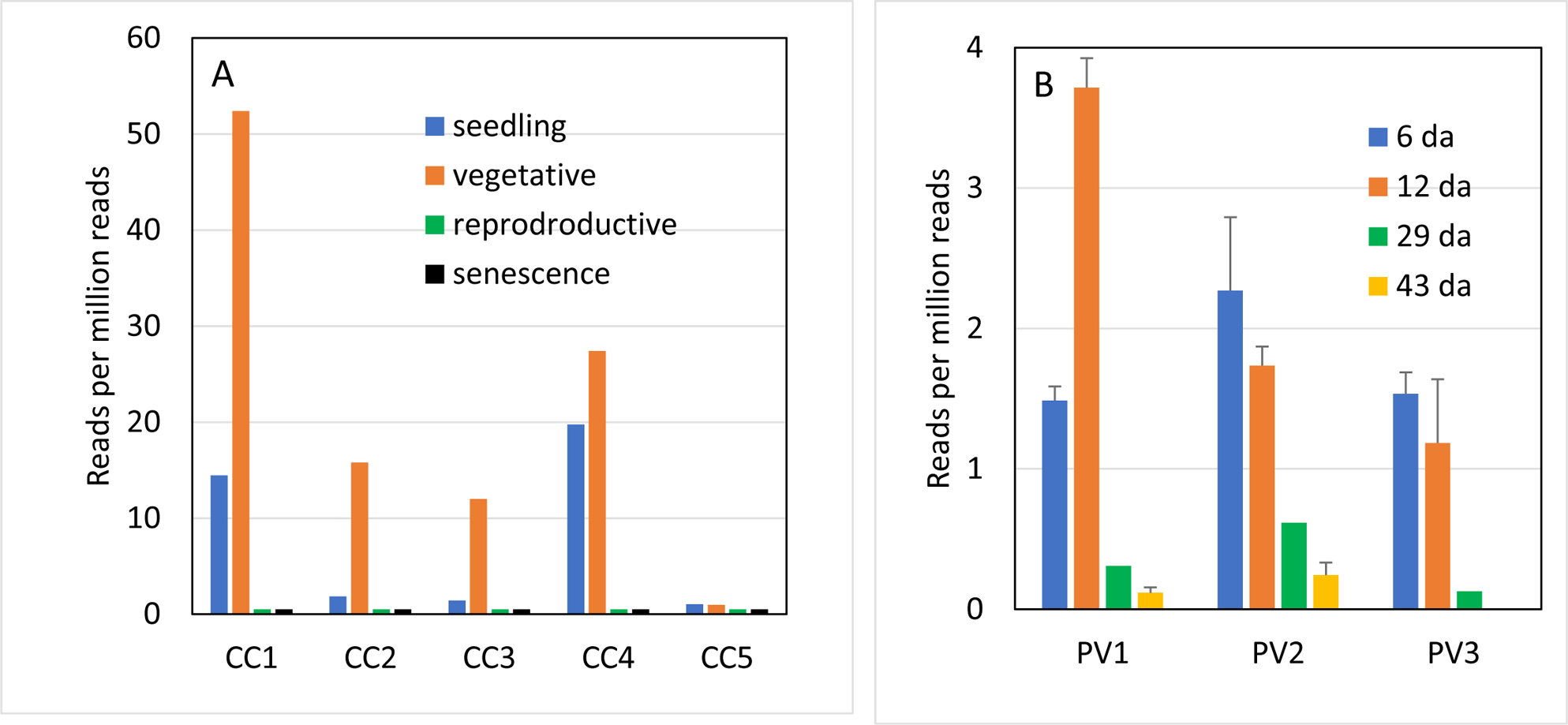
Expression of RCP genes in the root transcriptomes of beans at different stages of development. A, Chickpea cv. ICPL 87119 (single transcriptomes). B, Common bean cv. Bat93. (12-Day data is a combination of 10- and 14-day transcriptomes). Error bars show the range of two transcriptome replicates.

### Cicer arientinum (chickpea)

Chickpea is highly susceptible to heat stress. BioProject PRJNA748749 (Kudapa *et al*., 2023) looked for the molecular mechanisms of heat stress tolerance in *Cicer arientinum*, another major species of chickpea, by comparing the root transcriptomes of heat-sensitive and heat-tolerant cultivars. Heat stress (a gradual increase of about 10 °C over 15 days) was applied at both the vegetative and reproductive stages. A blast search found four RCPs for *C. arientinum* (Table S1). The RCPs were generally weakly expressed except for CA1, which was most strongly expressed in the transcriptomes of four of the cultivars in the vegetative stage. In the vegetative stage, its expression appeared to be downregulated by heat stress in three of the cultivars and upregulated in the fourth (Fig. 16). The up- and down-regulation did not appear to be associated with the heat sensitivities of the cultivars.

**Fig. 16.**
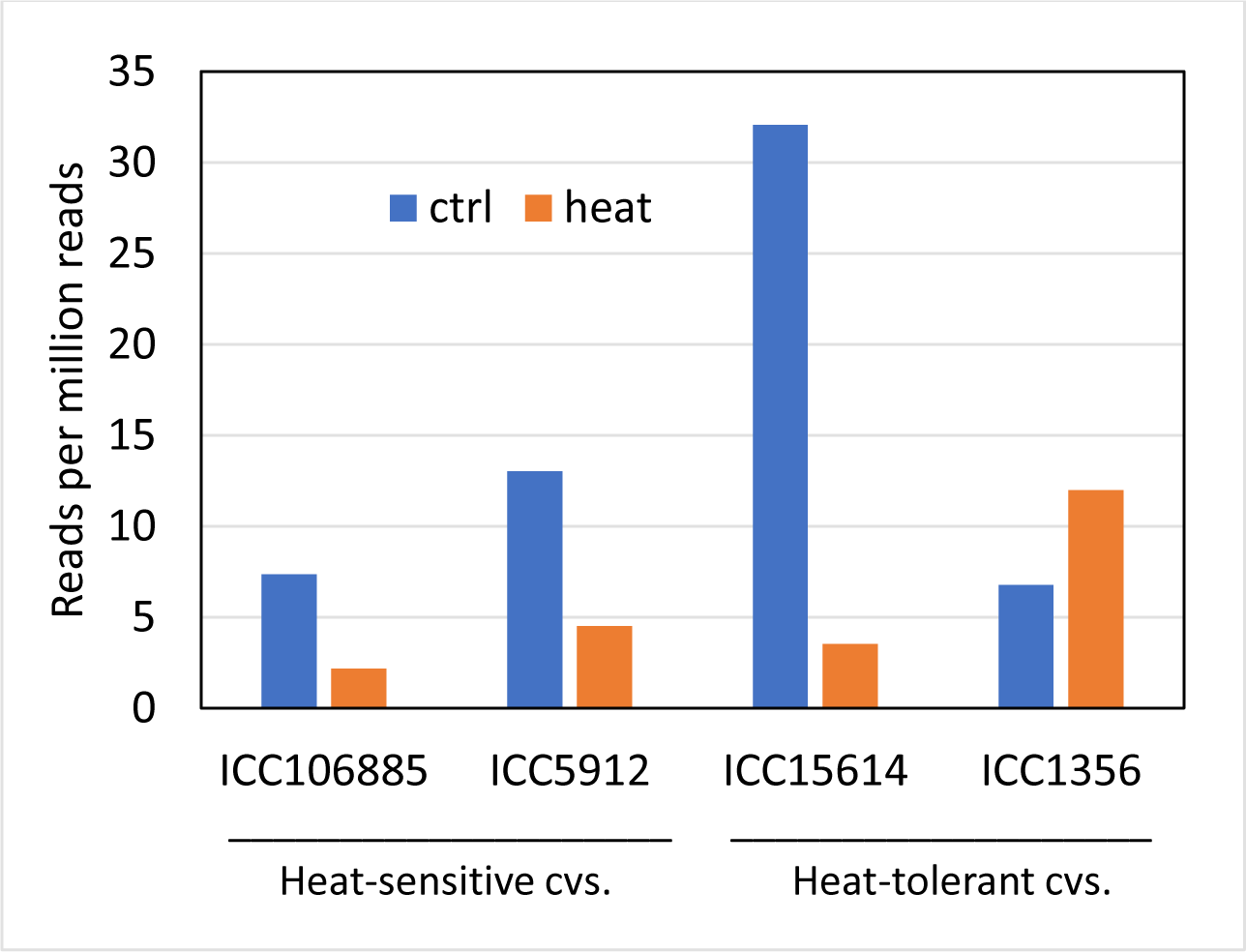
Effect of heat stress on the expression of RCP CA1 in the root transcriptomes of two heat-sensitive and two heat-tolerant chickpea cultivars at the vegetative stage.

### Cucumis sativa (cucumber)

Arbuscular mycorrhizal fungi (AMFs) have mutualistic relationships with most plant species. In cucumber, they have been shown to mitigate the effects of cold stress (Diagne *et al*., 2020). BioProject PRJNA316747 (Ma *et al*., 2018) examined the effects of an AMF (*R. irregularis*) and cold stress (14 days at 15 °C vs. 25 °C) on the root transcriptome of cucumber (cv. Zhongnong No. 26). A blast search found four RCPs (CS1-4) in the cucumber genome (Table 1). All four RCPs appeared to be strongly upregulated by AMF under both unstressed (Fig.17A) and cold-stressed (Fig. 17B) conditions. (However, AMF appeared to downregulate RCPs in rice (Fig. 14).

**Fig. 17.**
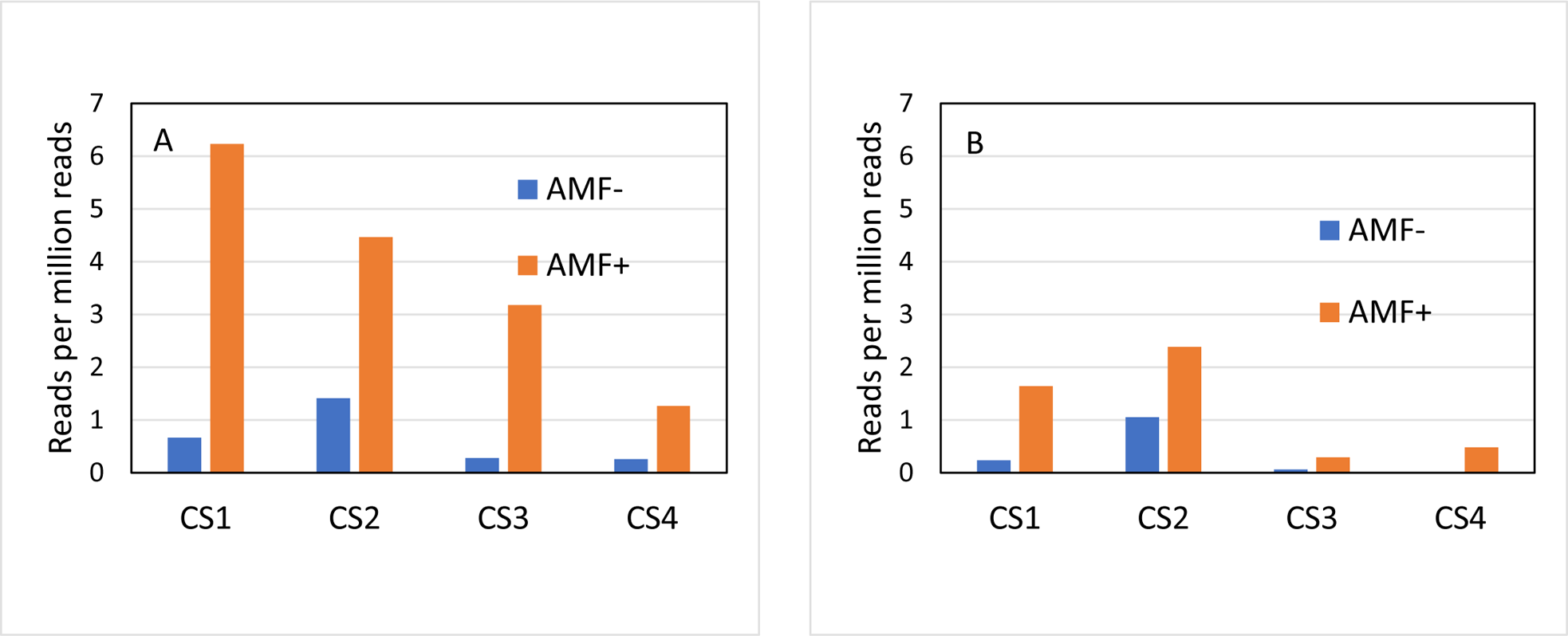
Effects of a mycorrhizal fungus on the expression of four RCP genes in the root transcriptomes of cucumber under normal and cold-stressed conditions. A, Control (14 days at 25 °C). B, Cold-stressed (14 days at 15 °C). One transcriptome was obtained for each condition.

## Discussion

Despite the abundance and ubiquity of RCP genes in land plants, it is surprising that they have received so little attention. Nothing is known of their functions other than that they are associated with the root cap, which has many roles in sensing and responding to the environment and controlling root growth (Arnaud et al., 2010). The first RCPs to be described were found in maize and one of them was shown to be expressed specifically in the outermost cells of the root tip that are continually being sloughed off into the soil (Matsuyama et al., 1999). These cells form part of a glycoprotein-rich mucilage that may help the root tip to penetrate the soil as well as bind to soil particles and improve communication with the soil (Matsuyama et al., 1999). Axenically produced maize mucilage was found to contain over 2800 proteins of many types (Ma *et al*., 2010), but it is unclear how many were RCPs.

The aim of this study was to see if RCP expressions in existing transcriptomes obtained under different conditions could provide some clues to their functions. This approach produced mixed results. Most of the transcriptomes examined did not show a clear response of RCP expression to the biotic and abiotic factors tested (Table S2). However, some RCPs appeared to be upregulated by stresses, including cold stress and lead stress in Arabidopsis (Fig. 2), low N in barley (Fig. 3A), salt stress (Fig. 4 B, C) and mycorrhizae disturbance (Fig. 5) in wheat, cold stress in rye (Fig. 6), maize (Fig. 7B) and sugarcane (Fig. 8) and salt stress (Figs. 10, 11) and low N and drought (Fig. 13) in rice. RCPs were also upregulated by mycorrhizal fungi in cucumber (Fig. 17) and during early developmental stages of beans (Fig. 15). On the other hand, other RCP appeared to be downregulated by the same stresses, including low N in a low N-tolerant barley cultivar (Fig. 3B), salt stress in a wheat cultivar (Fig. 4A), drought (Fig. 9), nematodes (Fig. 12) and mycorrhizal fungi and salt stress (Fig. 13) in rice and heat stress in chickpea (Fig. 15). RCPs also responded differently to the same stress in different cultivars of the same species, such as low N in barley (Fig. 3), salt stress in wheat (Fig. 4) and rice (Figs. 9, 10, 11) and low N in rice (Fig. 13). RCPs may not have central roles in these responses, as changing environmental conditions typically result in the up- and down-regulation of hundreds of genes. However, they may provide clues to designing further experiments to identify their functions.

In beans, RCP expression was strongest during early root development (Fig. 6) when roots are first adjusting to soil conditions and much signaling is expected to be occurring. However, their functions during this stage are unknown.

### Similarities to other protein families

As stated above, RCPs share several properties with small heat shock proteins. sHSPs are found in all kingdoms but are especially diversified in plants (Waters *et al*., 1996). They are best known for their strong upregulation by heat stress (Waters et al., 1996)). However, the present results do not support a similar role for RCPs as heat stress appeared to have mostly negative effects on RCP expression in maize (Figs. 7C) and chickpea (Fig. 16).

It was noticed that some RCPs were described as late embryogenesis abundant (LEA) proteins. LEA proteins are expressed in seeds during periods of drought and play a key role in the tolerance of plants to desiccation (Hernández-Sánchez *et al*., 2022). To see the degree of overlap between LEA proteins and RCPs, GenBank was searched for LEA proteins in the species examined in this study. Maize cultivar B73 was found to have 122 LEA proteins, seven of which were RCPs, and *Arabidopsis thaliana* cv. Columbia has about 14 LEA proteins, one of which was an RCP (Table S3). Only a few LEA proteins were found in the other species and none of them were RCPs. Zan et al. (Zan *et al*., 2020) classified 281 wheat LEAs by protein family and none of them were in the RCP family (PF06830). These results suggest that RCPs have only a weak connection to LEA proteins.

In addition to their similarities to small heat shock proteins outlined above, RCPs share a number of features with plant lectins, which might provide some clues to their functions. Lectins reversibly bind to carbohydrates with high specificity. In plants, they have roles in defense and interacting with symbiotic microbes (De Hoff *et al*., 2009) as well as in mitigating various abiotic stresses including temperature shock, drought and salt stress (for references, see (Jiang *et al*., 2010)). Like RCPs, plant lectin structures are abundant and diverse, have structures that are dominated by antiparallel beta sheets, are localized to roots and often have N-terminal secretion signals. Lectins, like small heat shock proteins (Waters et al., 1996), also tend to form multimers, which appears to be a property of beta-sandwich structures. It is thus possible that the β sheets of RCPs also interact to form multimers.

The N-terminal portions of RCPs are highly variable and have little homology to proteins of known function. A prominent exception is the ice-binding protein of the glacier-inhabiting *Ancylonema nordenskioeldii* (Procházková et al., 2024), which has a well-defined function. Another exception is TA9 (XP_044453695) of *Triticum aestivum*, whose N-terminal portion is shared by about 100 other land plant RCPs. AlphaFold was unable to model its structure with confidence.

### Origin of RCPs

*A. nordenskioeldii* is a chain-forming single-celled streptophyte alga and a member of the Zygnematophyceae, the closest known relatives of all land plants (Cheng *et al*., 2019). Its RCP, which includes the N-terminal secretory signal and C-terminal root cap domain of land plant RCPs, is an indication that RCPs developed early in the evolution of streptophytes, i.e., before the development of roots, which occurred approximately 400 mya (Raven & Edwards, 2001). Furthermore, a blast search shows that *Closterium*, a closely related alga, has about 60 RCPs (also with the characteristic N-terminal and C-terminal regions). This is a further indication of their importance well before the development of roots. Their functions remain unknown, but it seem likely that they, like the RCPs of higher plants, have roles in sensing the environment. The β-sandwich portion of RCPs appears to go back even further as it is found in a few other microorganisms, including a slime mold, a few bacteria and the primitive red alga *Porphyra*. In a tree of the β-sandwich regions, these organisms cluster separately from the streptophytes (Fig. 18). Learning the functions of these early proteins should provide valuable insights into the evolution of RCPs.

**Fig. 18.**
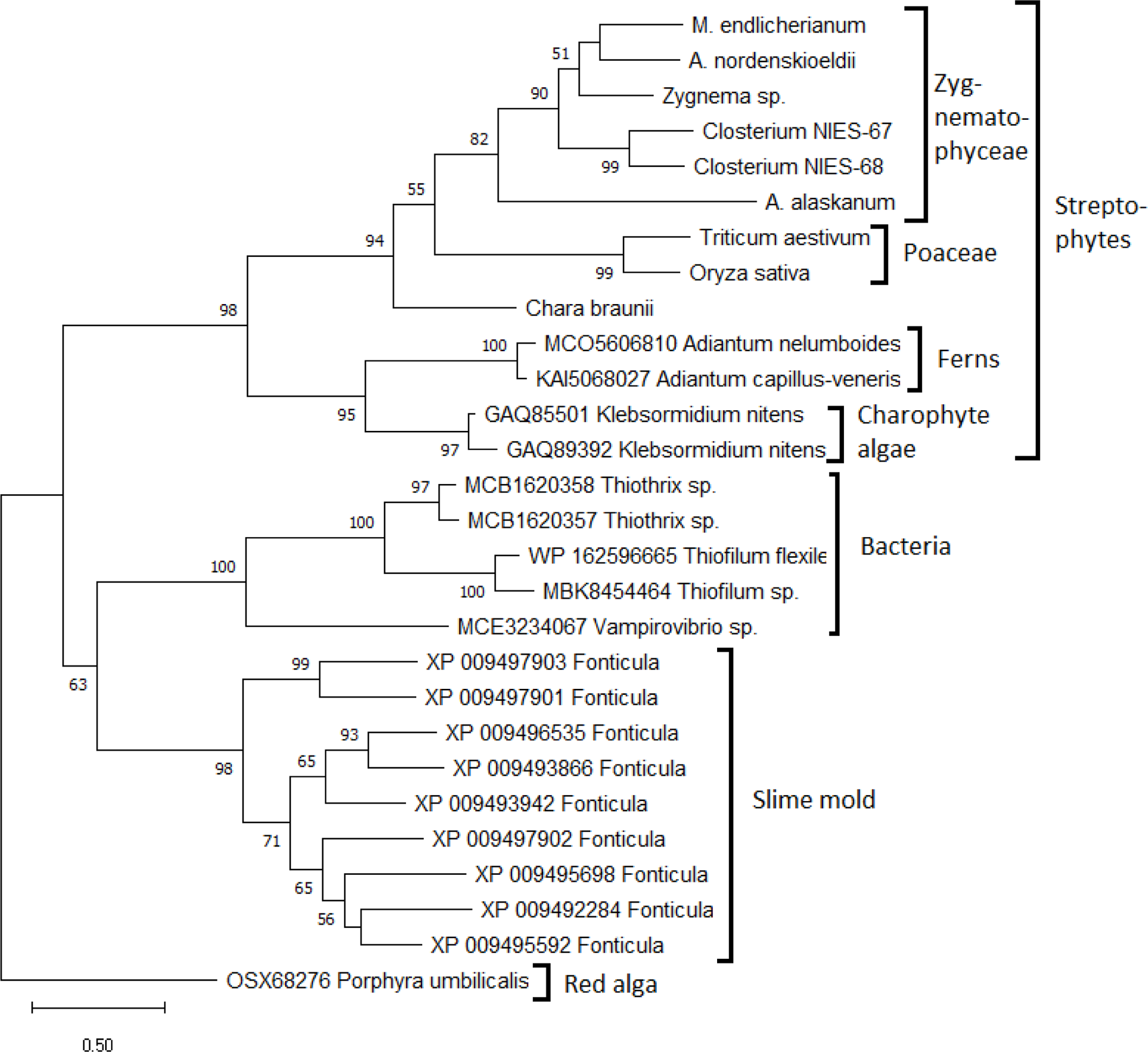
Maximum likelihood tree of the β-sandwich (DPR and DPH) domains in streptophyte RCPs and other organisms. Bootstrap values below 50% are not shown.

## Conclusions

The preceding results bring into focus a large family of plant proteins that have been largely neglected in the literature. The root cap of plant roots has been the subject of many studies because of its importance in root growth and development, but almost nothing is known about the root cap proteins, i.e., PF06830, that carry out many if not most of the functions of the root cap. The present results are a first step at understanding the functions of these proteins. They provide some hints at what their roles might be but don’t reveal any smoking guns. However, they should aid in the design of further studies to understand the roles and evolution of RCPs, as well as increase awareness of their existence, which they deserve.

## Acknowledgments

It is a pleasure to think Daniel Remias and Lenka Prochazkova for inviting me to join their investigations of *Ancylonema nordenskioeldii*, which lead to the present study. I thank the School of Life Sciences, University of Nevada Las Vegas for providing laboratory facilities for this study.

